# Native mass spectrometry of membrane protein lipid and ligand complexes using *de novo* designed WRAP domains

**DOI:** 10.64898/2026.07.24.740486

**Authors:** Surabhi Kokane, Ljubica Mihaljević, Barbara Abreu, Stijn de Schepper, Alexander Stevens, Erik G. Marklund, David Baker, Phillip J. Stansfeld, Michael Landreh

## Abstract

Membrane proteins engage in dynamic transient interactions with surrounding lipids along with their substrates. Capturing these concomitant contacts often remains a challenging problem. Here, we demonstrate that generative deep learning-designed WRAP domains can preserve weak interactions of membrane protein complexes in native mass spectrometry. Using WRAP-fused GlpG, AqpM, and OmpA as model systems, we show that lipid interactions can be retained and characterized without detergent micelles. We use the approach to show that WRAP-OmpA selectively binds phosphatidylethanolamine lipids within cavities formed at the protein–WRAP interface, while simultaneously accommodating weak chitobiose ligand binding. These findings establish WRAPs as versatile vehicles to probe ligand and lipid interactions of membrane proteins.

## INTRODUCTION

Membrane proteins generally exist in a complex environment, embedded in the lipid bilayer with hydrophilic surfaces exposed to the cytoplasm and cell exterior. Within the bilayer, the protein surface is coated with a shell of loosely associated annular lipids, which are in dynamic exchange with the surrounding bulk lipids, along with a distinct set of non-annular lipids that bind at specific sites.^1,2^ As a result, they often engage in multiple simultaneous interactions with the lipids in the bilayer and their surface-exposed ligands or substrates. Capturing these interactions has become increasingly important in recent years for understanding the mechanisms underlying the folding, oligomerization, and function of membrane proteins.

Native mass spectrometry has emerged as a useful tool to capture membrane protein-lipid complexes.^3^ Membrane proteins are ionized while embedded in detergent micelles or other membrane mimetics, or inserted into lipid vesicles.^4–6^ This protective environment is then removed through controlled collisions with gas molecules inside the vacuum region of the mass spectrometer to release the intact complexes for mass analysis.6–9 Soft-landing approaches have demonstrated that the tertiary and quaternary structures of membrane proteins remain essentially intact throughout the process.^10^ Similarly, tightly bound lipids or ligands remain attached during complex release.^11–13^ However, weak interactions, *e*.*g*. with annular lipids or labile ligands, are more sensitive to collisional activation and may therefore be distorted or lost during micelle or vesicle removal.^14^ As a result, the detection of labile ligand interactions has remained an open challenge in native MS.

As a possible solution, we turned to protein design. Machine learning has recently been used to design a solubilizing protein domain (WRAPs) around the membrane-facing region of the target membrane protein.^15^ The WRAP sequence is fused to its target membrane protein and provides a stably folded cage around the transmembrane region (TM). The inner surface of the WRAP domain is sterically and hydrophobically matched to the respective membrane protein, whereas the outer surface is characterized by a hydrophilic salt bridge network that ensures high stability and solubility. WRAPed membrane proteins are not membrane-inserted during translation and can readily be purified as soluble proteins.^15^ We therefore speculated that WRAPs may enable the preservation of labile ligand interactions with membrane proteins for native MS.

## RESULTS

### WRAPs can transport membrane proteins from solution to the gas phase

First, we evaluated the use of WRAPs as vehicles for membrane proteins in native MS. As target, we selected the *E. coli* membrane protease GlpG, which has been characterized extensively by native MS in detergent-solubilized form.^16,17^ GlpG acquires a high number of charges when released from DDM micelles at high activation energy (trap voltage 200 V), indicating partial un-folding.

*(Figure 1a*).^18^ Importantly, GlpG still retains a cardiolipin (CDL) molecule after detergent extraction, indicating a specific, functional lipid interaction.^16^

**Figure 1:**
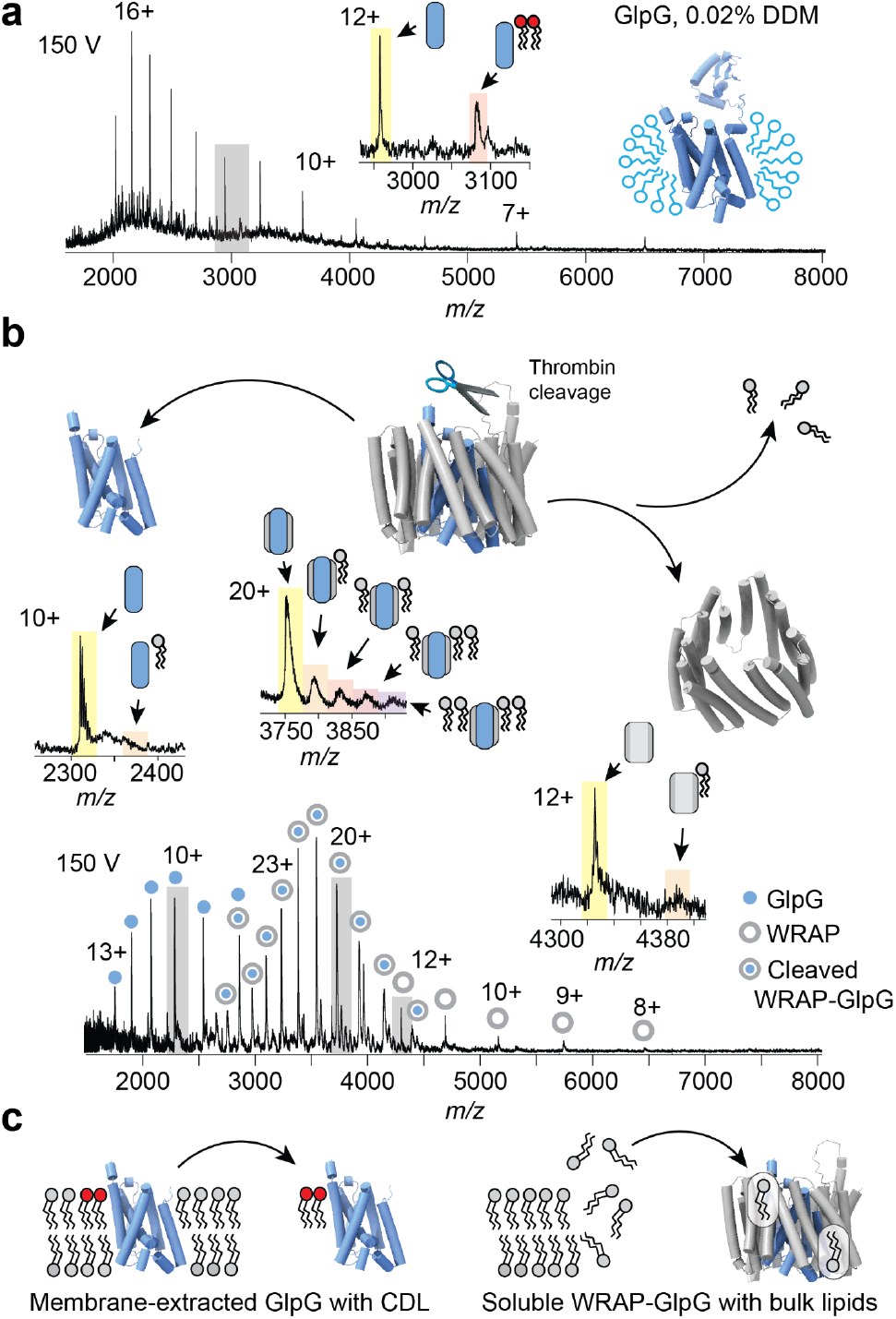
WRAPed GlpG binds membrane lipids. (a) Native mass spectrum of full-length GlpG released from DDM micelles at a trap voltage of 150 V shows highly charged protein ions and a specifically retained CDL adduct. (b) Mass spectra of the cleaved fusion protein show adducts consistent with lipid binding to the non-covalent GlpG-WRAP complex. Dissociation of the complex releases highly charged GlpG and lowly charged WRAP, and concomitant loss of lipid adducts. Charge states highlighted with gray shading are shown as zoom in the inserts. The predicted structure is shown as an insert with GlpG in light blue and the WRAP domain in grey. (c) Membrane-extracted GlpG retains CDL bound in the membrane, whereas WRAP-GlpG traps bulk lipids solubilized during purification.

We then subjected GlpG fused to an N-terminal WRAP domain via a polyglycine linker with a thrombin cleavage site to native MS analysis. The fusion protein could be analyzed without detergent and produced a single narrow charge state distribution which was sensitive to chemical unfolding, as expected for a soluble, globular protein (*Figure S1a, b*). Next, we performed enzymatic cleavage of the linker between WRAP and GlpG. Mass spectrometric analysis of the enzymatically cleaved WRAP-GlpG protein at an activation energy of 100 V showed mainly the intact WRAP-GlpG complex (*Figure S1c*). Raising the activation energy to 150 V, which is commonly used for detergent removal, resulted in three distinct ion populations, corresponding in mass to the intact fusion protein, free GlpG, and the empty WRAP domain. At 200 V, we observed loss of signal for the fusion protein, and increased signal for highly charged GlpG and lowly charged WRAP (*Figure S1c*). We conclude that GlpG remains non-covalently bound by WRAP after cleavage and can be dissociated and unfolded in the gas phase, analogous to the release of a native membrane protein from a detergent micelle (*Figure S1d*). Interestingly, we observed that the non-covalent WRAP-GlpG complex retains a low amount of adducts with an average size of 730 Da, in good agreement with the average mass of an *E. coli* membrane lipid, although we did not detect any adducts with a mass of 1.4 kDa that would indicate retention of CDL (*Figure 1b*) Interestingly, lipids are not observed for free GlpG or empty WRAP, suggesting that they might only interact with the intact GlpG-WRAP protein. WRAP-GlpG is not inserted into the membrane, making specific lipid retention unlikely. We speculate that the low-intensity adducts are annular lipids that are solubilized when bacteria are lysed during purification and trapped by the GlpG-WRAP fusion protein (*Figure 1c*).

### Lipids bind to membrane proteins inside the WRAP

Having established that WRAPs can transport and preserve membrane proteins in the gas phase, we further investigated their ability to interact with lipids. For this purpose, we selected Aquaporin M from *Methanoth-ermobacter marburgensis* (AqpM) and Outer membrane protein A from *E. coli* (OmpA) fused to N-terminal WRAP domains. Both proteins could be purified from the soluble fraction of *E. coli* expression cultures and buffer-exchanged into ammonium acetate in the absence of detergent. Native WRAP-AqpM and WRAP-OmpA can be detected without a need for extensive activation. Mass spectra show that both proteins exhibit up to four high-intensity adducts of ca. 730 Da each, consistent with the presence of co-purified lipids (*Figure 2a*).^15^ Lipid binding does not differ notably between charge states, unlike what has been reported for protein-detergent systems, which may be affected by charge redistribution upon detergent dissociation.^18,19^ We speculate that the even distribution of charged residues on the WRAP surface promotes stable gas-phase charge configurations that prevent lipid dissociation (*Figure S2a*).

**Figure 2:**
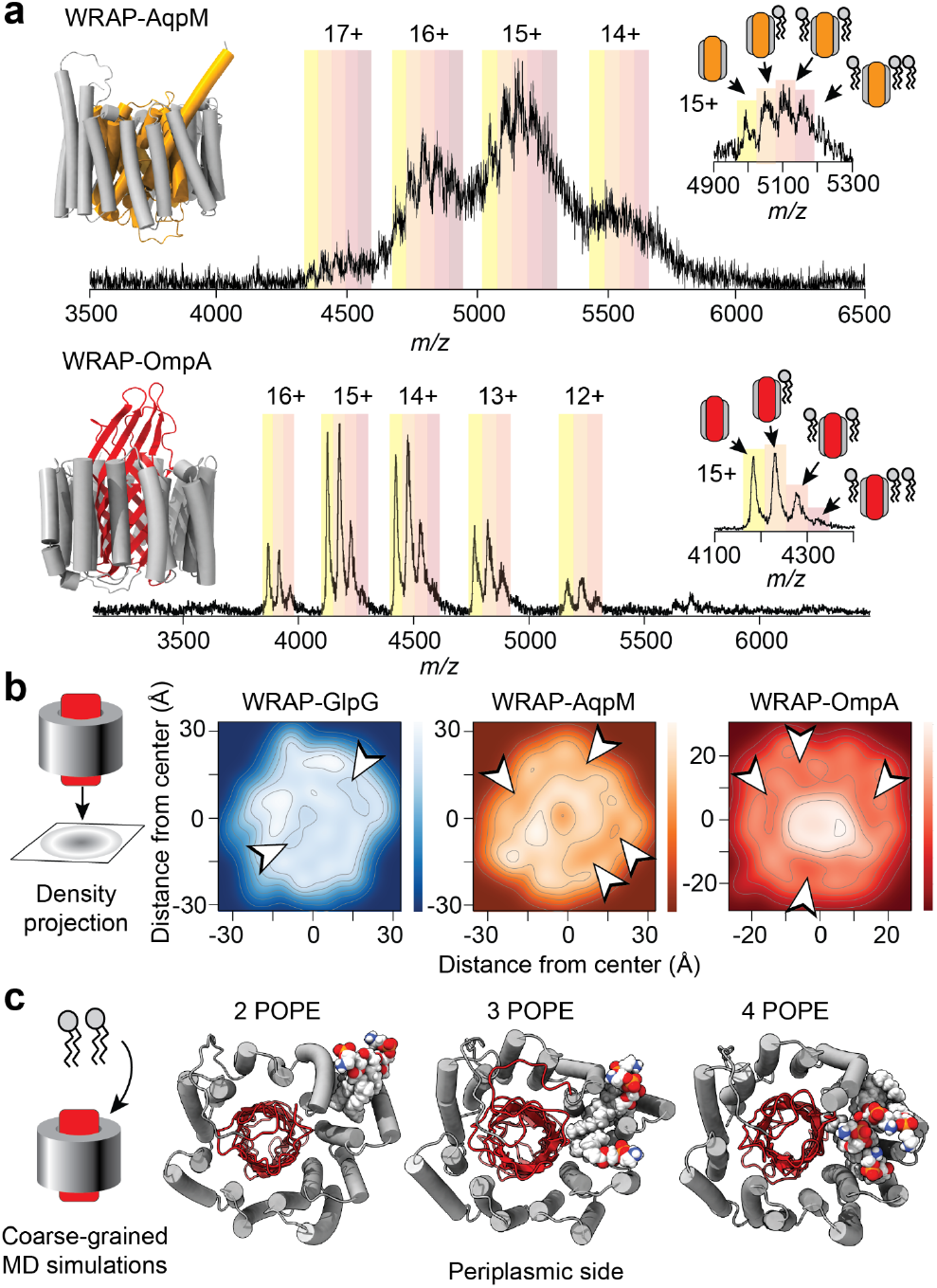
WRAPed membrane proteins bind lipids in native-like positions. (a) Native mass spectra of WRAP-AqpM (top) WRAP-OmpA (bottom) show prominent additional peaks corresponding to up to three adducts. The predicted structures are shown as inserts with the WRAP domains in grey. (b) Atom density projections along the transmembrane axis of WRAP-GlpG, WRAP-AqpM, and WRAP-OmpA indicate lose packing between the WRAP domain and AqpM or OmpA, but to a lesser extent for GlpG. Arrows indicate areas with low atom density. (c) CG-MD simulations of WRAP-OmpA with POPE clusters in solution show insertion of the lipids on the periplasmic side of the protein. Representative snapshots of simulations with two, three, or four POPE molecules are shown. WRAP is colored in gray, OmpA in red, and lipids are rendered as spacefill.

Dissociating lipids from detergent-solubilized membrane protein complexes usually requires an additional activation step.^20^ WRAPed protein complexes can be dissociated directly in the ion trap. Indeed, all adducts from WRAP-OmpA could be completely stripped off at an activation energy of 150 V (*Figure S2b*). To identify the adducts, we isolated the 13+ ion with a single adduct and recorded MS/MS spectra at a trap voltage of 100 V. Dissociating the complex resulted in multiple singly-charged peaks in the low *m/z* range consistent with sodiated phosphatidylethanolamine (PE) with acyl chains of 16 to 18 carbons (*Figure S2c*). Sodium adduction to the dissociated lipids may be promoted by the high density of acidic residues on the WARP domain and the WRAP-OmpA interface (*Figure S2b*) To test whether the lipids may be wedged between WRAP and the TM of OmpA, we employed an MS-based detergent competition assay.^21^ Briefly, the membrane protein-lipid complex is exposed to high concentrations of the detergent nonyl glucoside (NG) and analyzed by native MS. Lipids in annular positions are easily displaced by the detergent and sequestered in the empty micelles, whereas lipids in protected positions, such as high-affinity binding sites, remain bound. Mass spectra of WRAP-OmpA incubated with 0.5% NG recorded under gentle conditions show broad peaks, as expected for detergent binding. When the excess detergent was removed through activation in the gas phase, the lipid adducts remained essentially unaffected (*Figure S3a, b*). We conclude that the co-purified PE lipids are not sensitive to detergent competition, consistent with binding inside the WRAP domain (*Figure S3c*).

The fact that WRAP-AqpM and WRAP-OmpA display more pronounced lipid adducts than WRAP-GlpG suggests that they differ in their ability to accommodate lipids inside their WRAPs. To investigate this possibility, we computed the atom densities of the proteins and plotted the results as a density projection along the membrane axis (*Figure 2b*). WRAP-GlpG shows a relatively even atom density. WRAP-AqpM and WRAP-OmpA show a decreased density between membrane protein and WRAP, indicating free space that can potentially be occupied by lipids. To confirm that lipid binding to wrapped membrane proteins is sterically feasible, we performed coarse-grained molecular dynamics simulations (CG-MD) of WRAP-GlpG and WRAP-OmpA in solution in the presence of two, three and four POPE molecules diffused in an aqueous medium (*Figure 2c*). These simulations show that both WRAP-OmpA and WRAP-GlpG can accommodate up to four POPE molecules able to bind in the interface between WRAP and the membrane protein. In all cases, lipids were found to insert transiently and interact with the protein *via* their acyl chains and headgroups. In most replicas of the WRAP-GlpG simulations, the POPE molecules bind near transmembrane helices 1 to 4 and orient towards the cytoplasm (*Figure S4a*). This site is able to accommodate up to four POPE molecules and has been previously identified as a cardiolipin binding site in GlpG.^16^ Superimposition of GlpG simulated in the membrane^16^ with the WRAP-GlpG shows good agreement between the poses of two POPE molecules and the native CDL ligand at the primary site (*Figure S4b*). Secondary binding sites for POPE have also been observed in a subset of replicas. These sites are located near transmembrane helices 5 and 6, and near helices 3 and 6, in the vicinity of the catalytic helix and facing the periplasm (*Figure S4a*).^22^

WRAP-OmpA and WRAP-AqpM are also able to accommodate multiple POPE lipids in CG-MD simulations. For WRAP-AqpM, lipid clusters insert on the periplasmic side, where the lipids predominantly interact with transmembrane helices 1 and 2 (*Figure S4 c*). For WRAP-OmpA, lipids in most replicas tend to be trapped within the WRAP helices 11 to 14 and not interact with the OmpA interface, adopting a random orientation instead (*Figure S4d*). However, in a subset of replicas, POPE molecules diffuse towards the WRAP-OmpA interface and bind to OmpA. Two POPE interaction sites can be identified: The primary site is located near the trans-membrane strands 5, 6 and 8 and can accommodate up to four POPE molecules (*Figure 2c*), and the secondary binding site, which is occupied less frequently by 1-2 POPE, is located near strands 1 - 4 (*Figure S4c*). The headgroups of lipids in the primary site face the periplasmic side of the protein, but do not interact with the periplasmic loops. Lipids rather bind to the trans-membrane strands, as would happen in a membrane. POPE binds more persistently to WRAP-AqpM and WRAP-OmpA than to WRAP-GlpG, with some interactions lasting for 60% of the simulation time. The binding site adjacent to the β-strands 5 and 6, matches the AlphaFold prediction of the WRAP-OmpA complex with POPE. Interestingly, residues in β-strand 6 were found to be involved in the dimerization interface of OmpA.^23^ Together, these data suggest that WRAP-AqpM and WRAP-OmpA can retain lipids in physiologically relevant positions.

### WRAPs enable detection of concomitant lipid-ligand interactions with MS

The fact that we can detect complexes between WRAPed membrane proteins and lipids under gentle MS conditions suggests that the system can be used to probe simultaneous ligand and membrane contacts. To explore this possibility, we selected chitobiose (Glc-NAcβ1-4GlcNAc), a disaccharide present on the Ecgp glycoprotein that is expressed by brain microvascular endothelial cells.24 By recognizing its chitobiose moieties, OmpA acts as a receptor for Ecgp and in this manner mediates *E. coli* invasion in bacterial meningitis.^25^ Computational simulations and *in vivo* studies have shown that loops 2 and 3 of OmpA bind 1-2 chitobiose molecules.^26^

We previously used native MS to show ligand binding to WRAPed proteins.^15^ In line with these observations, addition of excess chitobiose to WRAP-OmpA resulted in several additional peaks corresponding in mass to protein-saccharide complexes (*Figure 3a*). However, chitobiose and lipid binding could not be distinguished directly, since two chitobiose molecules are similar in mass to a phospholipid. To determine complex stoichiometries, we took advantage of the fact that sugars bind relatively weakly in the gas phase whereas lipids require significant activation.^20,27,28^ Collisional activation of the WRAP-OmpA-chitobiose complexes revealed that the second and fourth adduct peak were removed near-completely at moderate trap voltages (60 - 70V). The third and fifth adduct peak, on the other hand, did not show a comparable reduction in signal intensity (*Figure 3a*). These observations allow us to assign the second peak as a bound chitobiose, the third peak as a bound lipid, and the fourth peak as a complex with both a chitobiose and a lipid molecule. Notably, the trap voltages that dissociate the chitobiose complexes are below the range commonly required to dissociate detergent micelles from membrane proteins.^7^ Therefore, these labile interactions would essentially be inaccessible without the use of WRAPs as membrane protein vehicles.

**Figure 3:**
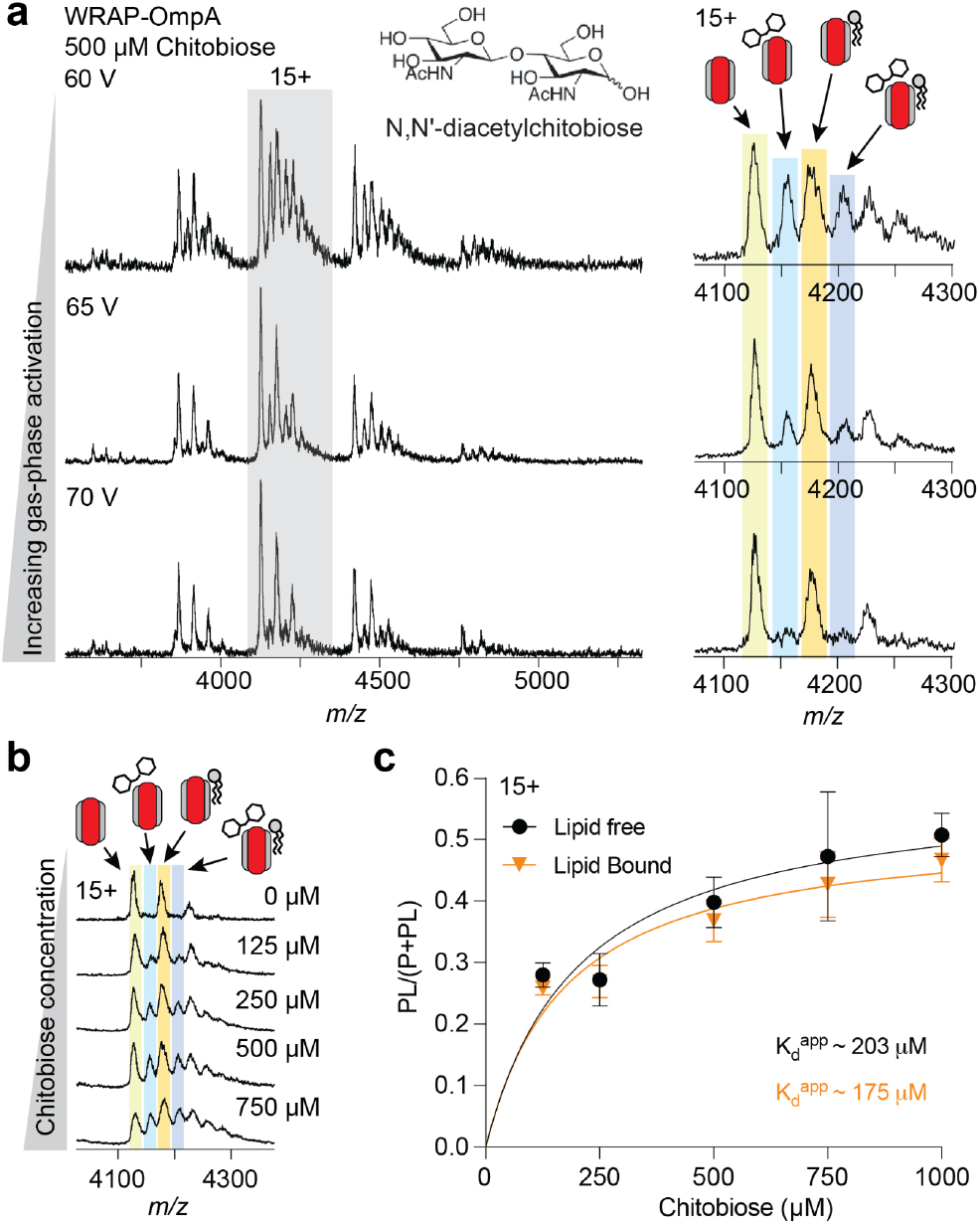
MS captures chitobiose binding to WRAP-OmpA-lipid complexes. (a) Mass spectra of WRAP-OmpA with chitobiose recorded at increasing trap voltages show facile dissociation of the saccharide ligand, but not the lipid adduct. A zoom of the 15+ charge state is shown to the right. (b) Zoom of the 15+ charge state at increasing chitobiose concentrations shows that increasing ligand binding can be monitored by native MS. (c) Plotting the ratio of PL to P + PL for WRAP-OmpA and WRAP-OmpA-PE binding to chitobiose for the 15+ charge state produces near-identical binding curves. Error bars indicate average and standard deviation (n = 4).

Since we can resolve simultaneous lipid and chitobiose binding, we asked whether both interactions affect each other. As outlined above, the co-purified PE molecules likely resemble native membrane interactions with regard to both lipid type and location, making the insights potentially valid in the biological system. Molecular docking of a chitobiose molecule into the predicted structure of the WRAP-OmpA-lipid complex confirms ligand binding to the disordered loops L1-L3.^26^ Plotting the fraction of intact chitobiose complexes with and without bound PE as a function of trap voltage shows essentially identical stabilities for the chitobiose complexes with WRAP-OmpA and with WRAP-OmpA plus one PE (*Figure S6a*). Next, we titrated WRAP-OmpA with chitobiose and monitored the change in signal intensities for the different ligand and lipid populations (*Figure 3b*). We then calculated the ratios of chitobiose-free and chitobiose-bound protein at each concentration. Plotting the PL/(PL+P) ratios as a function of ligand concentration shows no significant difference between chitobiose binding to WRAP-OmpA or to WRAP-OmpA with one PE adduct (*Figure 3c, Figure S6b*).

The highly similar stabilities and binding curves suggest that PE does not affect chitobiose binding. To contextualize this finding, we considered the native topology of OmpA. Chitobiose binds to the extracellular side of OmpA in the outer *E. coli* membrane during pathogen-host contacts. The outer leaflet is composed mainly of lipopolysaccharides, whereas the inner leaflet is rich in PE, which interacts non-specifically with the periplasmic side of OmpA.^29^ Consistent with the lipid distribution in the outer membrane, our CG-MD simulations show spontaneous PE insertion on the periplasmic side of the WRAP complex (*Figure 2c*). Since these lipids are therefore far removed from the chitobiose binding site on the extracellular side, they are unlikely to affect ligand binding (*Figure S5c*).

## CONCLUSIONS

Our results highlight the potential of WRAPed membrane proteins in simultaneous detection of lipid and ligand interactions under gentle native mass spectrometry conditions. By stabilizing transient contacts that are otherwise lost during conventional sample preparation, WRAPs provide new opportunities to dissect the interplay between membrane composition, protein folding, and functional ligand recognition. Future work may focus on rationally designing WRAP architectures with tailored lipid pockets or dynamic interfaces to better mimic native bilayers. Integrating this strategy with computational modelling, structural MS workflows, and functional assays could expand its applicability to diverse membrane protein classes, facilitating mechanistic insight and aiding drug discovery efforts.

## ASSOCIATED CONTENT

### Supporting Information

Experimental Methods, Figure S1: Dissociation of WRAP-GlpG, Figure S2: Dissociation of WRAP-OmpA-lipid complexes, Figure S3: Detergent titration of WRAP-OmpA, Figure S4: MD simulations of WRAP-lipid complexes, Figure S5: Dissociation of WRAP-OmpA-ligand complexes. The Supporting Information is available free of charge on the ACS Publications website.

## AUTHOR INFORMATION

## Author Contributions

SK and ML designed the study with input from DB and PJS. SK produced proteins. SK and AS conducted mass spectrometry measurements. LM designed WRAPs and provided protocols. BA performed CG simulations under supervision of PJS. SdS performed density calculations under supervision of EGM.

