## Supplementary File for "Native mass spectrometry of membrane protein lipid and ligand complexes using *de novo* designed WRAP domains"

### Methods

#### Protein design and production

Full-length GlpG was expressed and produced as described.[1] WRAPs for GlpG, AqpM, and OmpA were designed previously. Synthetic genes fusions of WRAP designs and target membrane protein (OmpA, GlpG and AqpM, Table S1) with flexible GS linker, thrombin cleavage site and C-terminal histidine tag were synthesized by Genscript Inc in pET11b expression vector. These plasmids were transformed into *E. coli* BL21 (DE3) (New England BioLabs) and plated on LB agar plates containing 100 µg/mL of Ampicilllin (Merck). For protein expression, the colonies were inoculated for overnight precultures at 37 °C and further added to 500 ml LB media for large scale expression. The cultures were incubated at 37 °C until they reached an OD600 of 0.5 and were later induced with 0.2 mM isopropylthio-β-galactoside (IPTG, Sigma). Post-induction, the cultures were incubated at 16 °C overnight for protein expression. Cells were harvested by centrifugation at 8000 x *g* for 15 mins. Cell pellets were homogenized in a buffer containing 20 mM Tris-HCl pH 8.0, 150 mM NaCl and supplemented with protease inhibitor cocktail (Thermo scientific). The cells were lysed using probe sonicator (5 min, 10s on, 5s off, 40%) and centrifuged at 10,000 x *g* for 10 min. The lysate was loaded onto a Ni-NTA affinity resin (Qiagen) and subsequently washed with buffers containing 20mM Tris-HCl pH 8.0, 150mM NaCl, 30 mM and 50 mM imidazole. The proteins were eluted in buffer containing 20mM Tris-HCl pH 8.0, 150mM NaCl, 250 mM imidazole and dialysed overnight with 3.5 kDA MWCO dialysis-tubing (Snake skin, Thermo scientific) in buffer containing 20mM Tris-HCl pH 8.0, 150mM NaCl. The sample was concentrated using a 30-kDa MW cutoff concentrator (4ml, Millipore) at 7000 × *g* and further purified using ÄKTA go (Cytiva) with Superose 6 increase 10/300 (Cytiva) column. The fractions with monodispersed peaks were pooled together and further used for native-MS.

#### Native mass spectrometry

WRAP-OmpA, WRAP-GlpG, and WRAP-miniCXCR4 were buffer-exchanged into 500 mM ammonium acetate, pH 7, milliQ, and 100 mM ammonium acetate, pH 7, respectively, using BioSpin6 columns (BioRad, Hercules, CA, USA). Wild-type GlpG was buffer-exchanged into 100 mM ammonium acetate supplemented with 0.02% DDM (Sigma). Samples were loaded into nESI capillaries (Thermo Fisher Scientific Inc., Waltham, MA, USA). Mass spectra were recorded in positive ionization mode on a Waters Synapt G1 traveling-wave IM mass spectrometer (MS Vision, Almere, The Netherlands). The capillary voltage was maintained at 1,5 kV and the sample cone was 100 V. The source temperature was 30°C. The transfer collision energy was 10 V and the trap voltage was adjusted between 10 V and 200 V as described in the text. The trap gas was argon at a flow rate of 4 mL/h, the IMS gas was nitrogen at a flow rate of 30 mL/h. Data were analyzed using the Waters MassLynx 4.1 and mMass V3.0 software packages.

**Coarse-grained POPE diffusion molecular dynamics simulations**

AlphaFold 3 was used to generate protein models of WRAP-OmpA, WRAP-AqpM, and WRAP-GlpG.[2] For the coarse-grained molecular dynamics (MD) simulations, the Martini 3 force field was used, along with the GROMACS 2024.5 simulation package.[3,4] Each structure was converted to coarse-grained resolution using Martinize2.[5] A force constant of 200 kJ/mol was used for the elastic network potential with a maximum distance cutoff of 0.7 nm.  The protein-WRAP was enclosed in a box, solvated and ionized with 0.15 M NaCl, with additional ions to keep system neutrality. A single POPE molecule was added to the initial systems. The simulations were performed at the temperature of 323K. An initial energy minimization was performed, followed by a first step of equilibration at constant volume for 5 ns, and a second step of equilibration at constant pressure for another 5 ns. During equilibration, the temperature was controlled using a velocity-rescale thermostat, with a Berendsen barostat for pressure control.[6,7] Thirty replicates were initially simulated for 200ns.  A maximum of five WRAP conformations with POPE bound were selected, and another POPE was added to the system, followed by a new solvation and ionization process. The same setup and equilibration process described above was used, and thirty new replicates with two POPE molecules were generated and simulated for 200 ns each. For the production simulations the C-rescale barostat was used instead.[7] Out of all the conformations with two POPE bound, a maximum of five conformations were selected to be used as seeds and starting points to study the binding of three POPE molecules using thirty replicas simulated for 200 ns each. In a similar way the same procedure was used to study the binding of four POPE to either GlpG or OmpA complexes. Final conformations from the MD simulations were converted to an all-atom resolution using the CG2AT package.[8]

**All-atom molecular dynamics simulation of GlpG-POPE, OmpA-POPE and OmpA-chitobiose complexes**

AlphaFold 3 was used to build the initial complex models with POPE, while the OmpA-chitobiose system was generated with SwissDock, because AlphaFold placed chitobiose near OmpA N-terminus contradicting the experimental evidence that suggests otherwise.[9,10] The forcefield used was CHARMM36.[11] CGENFF was used to parameterize chitobiose, while the POPE parameters used were the ones already contained in the CHARMM forcefield. The systems were prepared for simulation using the tools on the GROMACS 2024.5 simulation package.[3] Each system was enclosed in a box, solvated and 0.15 M NaCl ions was added, alongside with extra ions to neutralize the system. Each system was minimized using steepest descent followed by three equilibration steps, one at constant volume for 100 ps with position restraints on the heavy atoms of the system, the second at constant pressure with position restraints for 500 ps, and the final one without position restraints for another 500 ps. The temperature was set at 310 K using a velocity rescale thermostat and the pressure was controlled using C-rescale barostat.[6,7] Five replicas of 200 ns each were done. The interactions between ligands and protein were analysed using MDAnalysis and renderings of atomistic models generated from the CG-MD data with CG2AT2 were done using ChimeraX.[8,12,13]

**Atom density calculations**

Calculations were performed using the MDAnalysis (version 2.10.0)[13] and scipy (version 1.16.3)[14] python packages. WRAP-OmpA, WRAP-GlpG and WRAP-AqpM structures are predicted by AlphaFold 3.0. The C-terminal tails (from residues 544 for OmpA, 654 for GlpG and AqpM) were excluded, to avoid interference in density calculations. Structures were aligned along the transmembrane axis. Densities were calculated by projecting atom positions on the perpendicular plain and the distribution was smoothed using a Gaussian Kernel Density Estimation.

**Table S1. Sequences of the WRAP fusion proteins used in this study.** WRAP sequences are underlined.[15]

| Protein Name | Sequence |
|---|---|
| WRAP-GlpG | MSGRERLDRLYLEVLELRAEVLELVQEGAPAEELVRAAERLVD<br>ATLELLDLEAEEAEDLMERATLLTMRAVVLAMRLFLAVGVGAPE<br>EEIERLVEELERVIEETEKLLEEIKGGVESLLTALALKILLSLVKSL<br>VLTLLGAPAEEIKEFTDKQLKELLELLRETIEKAREAGAQGELFE<br>ARVLLTGLEVMKKLRETGSVLPSDELVFELVSGILRALLETAARV<br>TGAEEEMRRLYEEAEERLRRGLAEIASLPAEEADEAFALLLAQV<br>LRDVLVAILELVDKRAAELVRALLDFIIALAETKLEVLKEKDEEEA<br>KKKALEGLEKVEELAKKLIELRFKDSPILENLKNAITYATELFKAL<br>VEGAELEEIEALFDKLAEYMDKAVELEVKSLPPVEAARLRHTAR<br>AGLALLRAAAAARAGRREEAEREALREFNRLVLEAARETLELE<br>RRADYQGGGGSGGGGSLVPRGSGGGGSGGGGSERAGPVT<br>WVMMIACVVVFIAMQILGDQEVMLWLAWPFDPTLKFEFWRYFT<br>HALMHFSLMHILFNLLWWWYLGGAVEKRLGSGKLIVITLISALLS<br>GYVQQKFSGPWFGGLSGVVYALMGYVWLRGERDPQSGIYLQ<br>RGLIIFALIWIVAGWFDLFGMSMANGAHIAGLAVGLAMAFVDSL<br>NAGSGSHHWGSTHHHHHH |

| | |
|---|---|
| WRAP-AqpM | MSGELEEAFKLITQLARDFFKTAAEIFREQGNELGAKIMEVYAK<br>YFEEMIKNPEEAPRLFLKMHADVARVIAEFARKRGDLAAAKMA<br>EAAAKFFELLAENPEEFQEAFLKGAALLMRVGAEEAREAGDEA<br>LAEALERAAELFEEALKDPSKAGEYLVEAMRVLAEALAELAEER<br>GDLATARALRTLVETLKEIEKNKDNLHLEVERLTLRLMLVNLRAA<br>AEDGWTEFLRVADILEKAIEALYGSEEDRKRYTLEVAAELARVV<br>SRYARRVGDELGARLFEVVAERLEKALETLDKIEEEMSKALVD<br>MLRAAAEWVRTQGNEDAAEILDLLAKIMESIINNPENFAKEAKE<br>YLEKIKDLSIKSLEKRGDTDGVKLFSVLSDSFIKLLEFEIEKDYQ<br>GDRSGGGGSGGGGSVSLTKRCIAEFIGTFILVFFGAGSAAVTL<br>MIASGGTSPNPFNIGIGLLGGLGDWVAIGLAFGFAIAASIYALGNI<br>SGCHINPAVTIGLWSVKKFPGREVVPYIIAQLLGAAFGSFIFLQC<br>AGIGAATVGGLGATAPFPGISYWQAMLAEVVGTFLLMITIMGIAV<br>DERAPKGFAGIIIGLTVAGIITTLGNISGSSLNPARTFGPYLNDMIF<br>AGTDLWNYYSIYVIGPIVGAVLAALTYQYLTSEGSRDRSGSGSH<br>HWGSTHHHHHH |
| WRAP-OmpA | MSGELEQKAVETIAEMAIQMMLGGPSEEIRELWDKFIALIEEITA<br>ATPVETALARAMAELVRNLALGKPIDEIEKSAREFAAKLKELYGD<br>RTAEAEAILAIVAAVKAGIEGKPKEEIAELLARYRDAVAKILADDP<br>ARQAVMTAVLDLIAALLLGEPAEEVRARLAELEALIDRLFDQLTP<br>EELMFAAYTLLLAALAIGLPAEELLALLERLVARLREALADDPEAL<br>HIAELHLRVLVLTQLGKPIEEILELIKEILEEYKEELGEPTASF<br>LVHALSLLLAVAISSRDTWELAKPLIDELLKGALEKLEDENVANYI<br>KAFADLMKAAIEGKPIEELAELFRKMTEYLIKIIDYQGGGGSGG<br>GGSLVPRGSGGGGSGGGGSAPKDNTWYTGAKLGFSQYHDT<br>GFINNNGPTHENQLGAGAFGGYQVNPYVGFEMGYDFLGRMP<br>YKGSVENGAYKAQGVQLTAKLGYPITDDLDIYTRLGGMVFRAD<br>TKSNVYGKNHDTGVSPVFAGGVEYAITPEIATRLEYQFTNNIGD<br>AHTIGTRPDNGMLSLGVSYRFGQGEAAGSGSHHWGSTHHHH<br>HH |

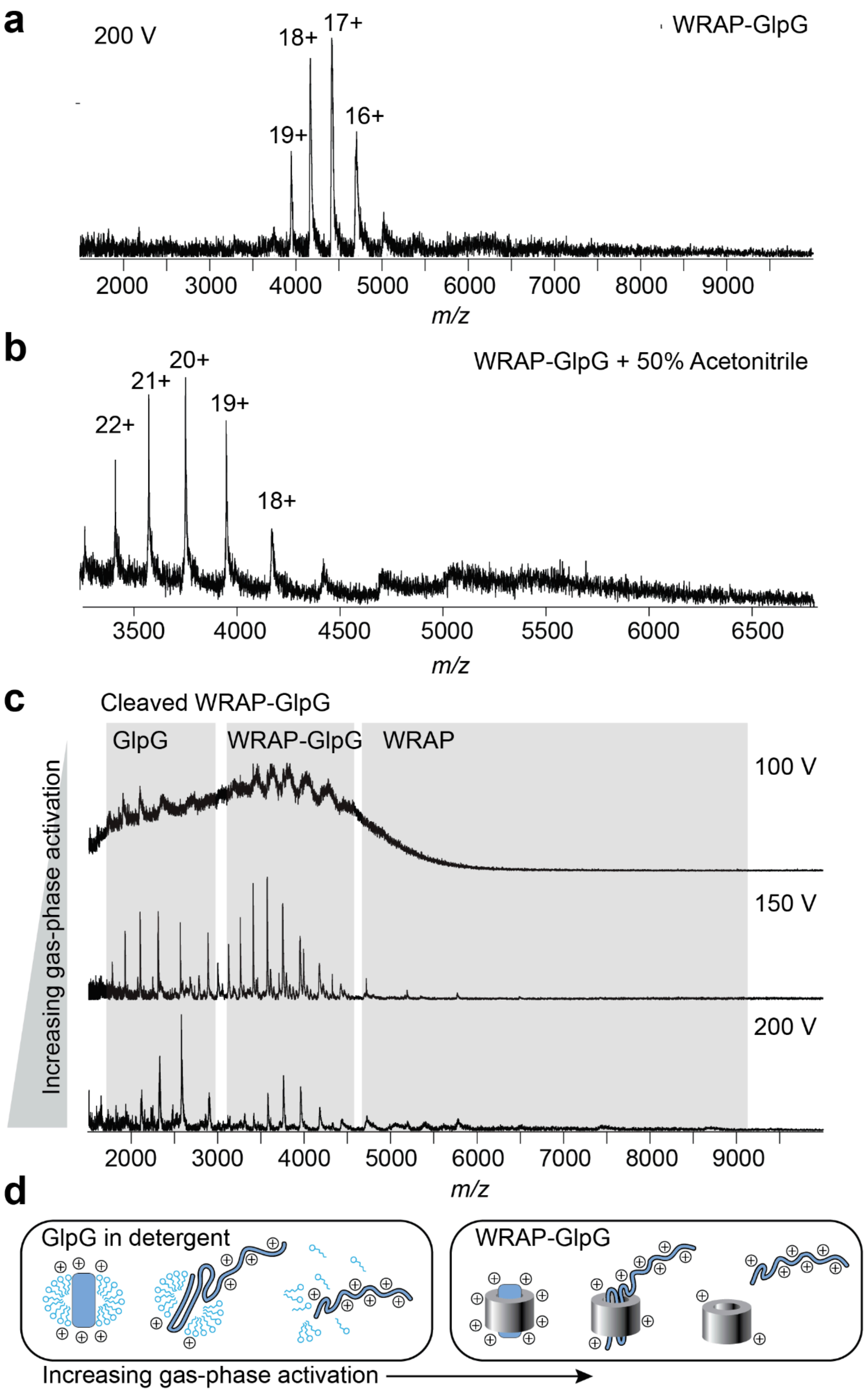


***Figure S1.*** *Mass spectra of intact (a) and denatured (b) WRAP-GlpG show a single charge state envelope and no dissociation even at high trap voltage. (c) Native MS analysis of thrombin-cleaved WRAP-GlpG causes dissociation of GlpG and the WRAP domain when the trap voltage is raised. (d) Proposed release of unfolded, highly charged GLpG from detergent micelles (left) and WRAP (right).*

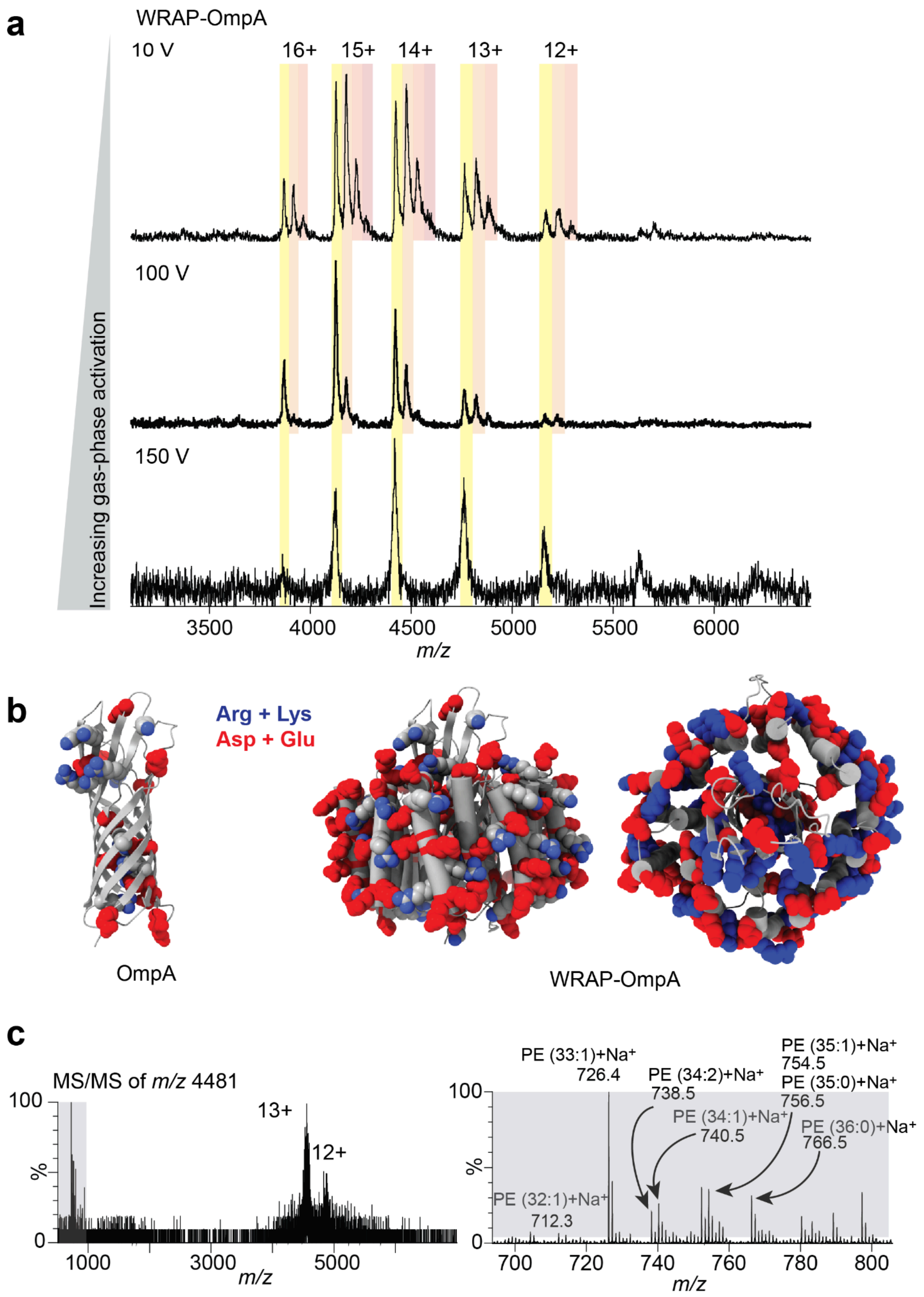


***Figure S2.*** *(a) Ionizable residues are sparse on the surface OmpA (left) but abundant on the surface of the WRAP domain of the WRAP-OmpA fusion protein (shown as side view and from the extracellular side, middle and right). (a) Native mass spectra of WRAP-OmpA show prominent additional peaks corresponding to up to three adducts which can be dissociated through gas-phase activation. (c) MS/MS spectra of the 13+ complex with one adduct results in dissociation of the adduct to release a 12+ apo peak and several low-molecular species (left). The low-molecular weight region of the MS/MS spectrum shows multiple peaks corresponding in mass to sodiated PE lipids with varying chain lengths.*

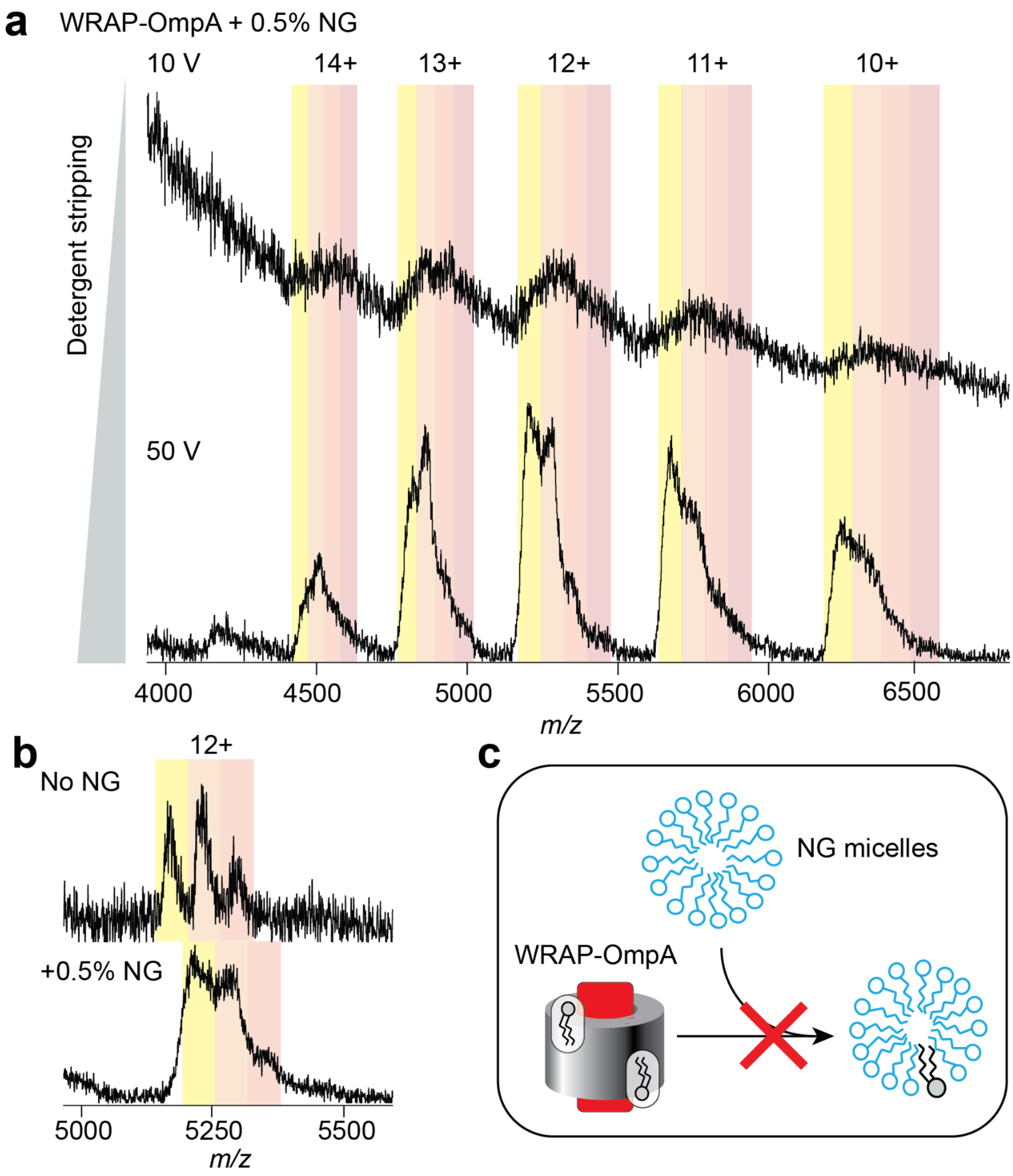


***Figure S3.*** *(a) Native mass spectra of WRAP-OmpA in the presence of 0.5% NG show peak broadening at low trap voltage (top, 10 V) due to detergent binding. At moderate trap voltage (bottom, 50 V), the detergent is partially dissociated and individual charge states of WRAP-OmpA can be resolved. (b) Comparison of the 12+ charge state of WRAP-OmpA without (top) and with NG (bottom) at a trap voltage of 50 V shows no major change in the intensity of the lipid adducts. (c) Schematic of the detergent competition MS assay. Lipid adducts on WRAP-OmpA are not removed in the presence of an excess of NG micelles, indicating binding to protected site(s).*

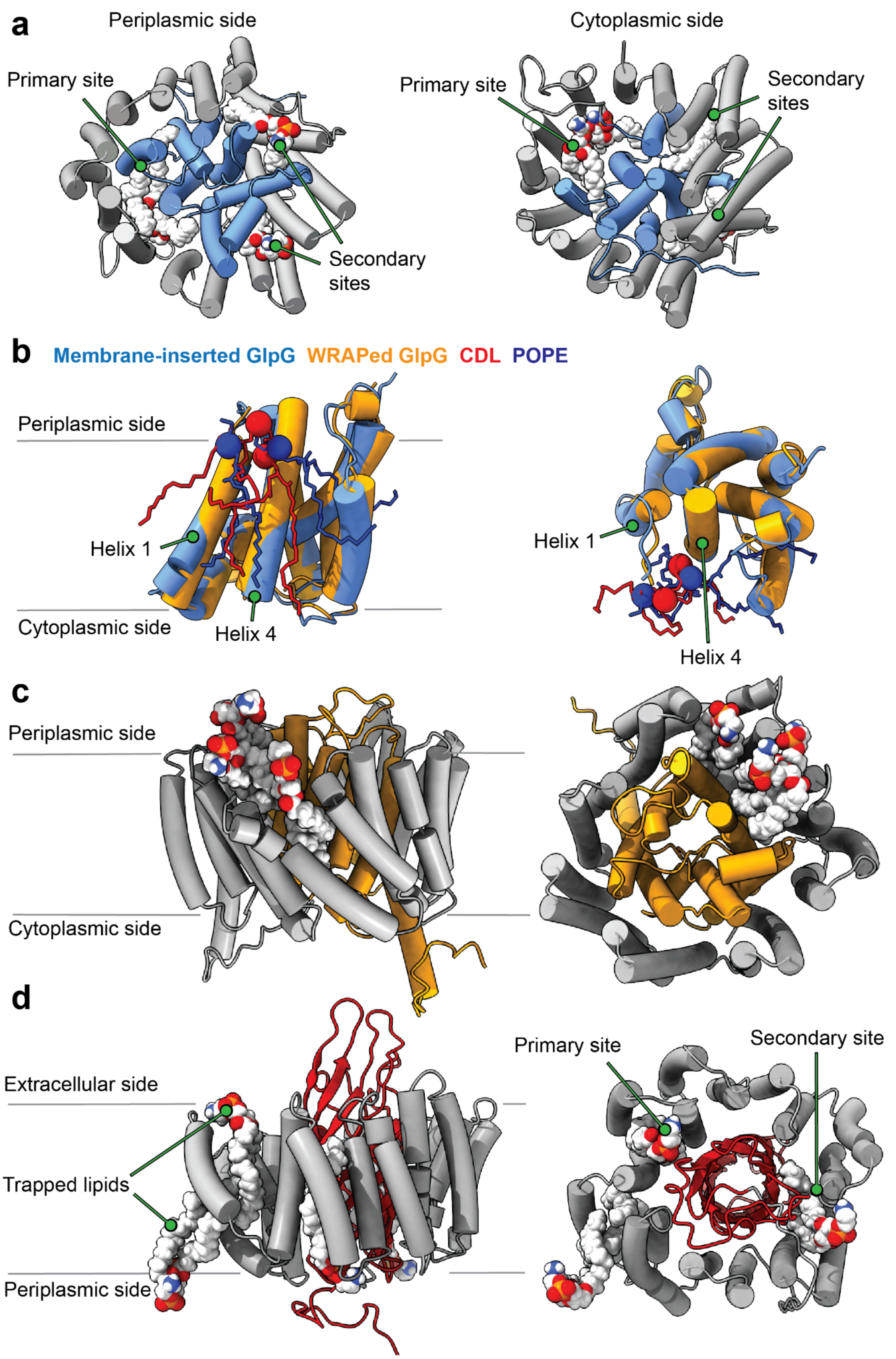


***Figure S4.*** *(a) Representative CG-MD snapshot converted to an atomistic model of WRAP-GlpG with POPE shows occupation of the primary lipid binding site on the cytoplasmic side of the protein, as well as two secondary sites with one POPE molecule each on the periplasmic side. Lipids are rendered as spacefill, the WRAP domain is colored grey and GlpG is colored blue. (b) Comparison of GlpG (blue) in*

*complex with CDL (red) simulated in the membrane[16] with WRAP-GlpG (orange) simulated with two POPE molecules (red) at the primary lipid binding site. The WRAP domain is omitted for clarity. The overlaid structures are shown as side view (left) and from the cytoplasmic side (right). Phosphate atoms in the lipid head groups are shown as spheres. (c) Representative CG-MD snapshot converted to an atomistic model of WRAP-AqpM with 3 POPE molecules show insertion on the periplasmic side of the protein. (d) Representative CG-MD snapshot converted to an atomistic model of WRAP-OmpA with POPE shows occupation of the primary as well as a secondary lipid binding site and lipids trapped in random orientations between the WRAP helices.*

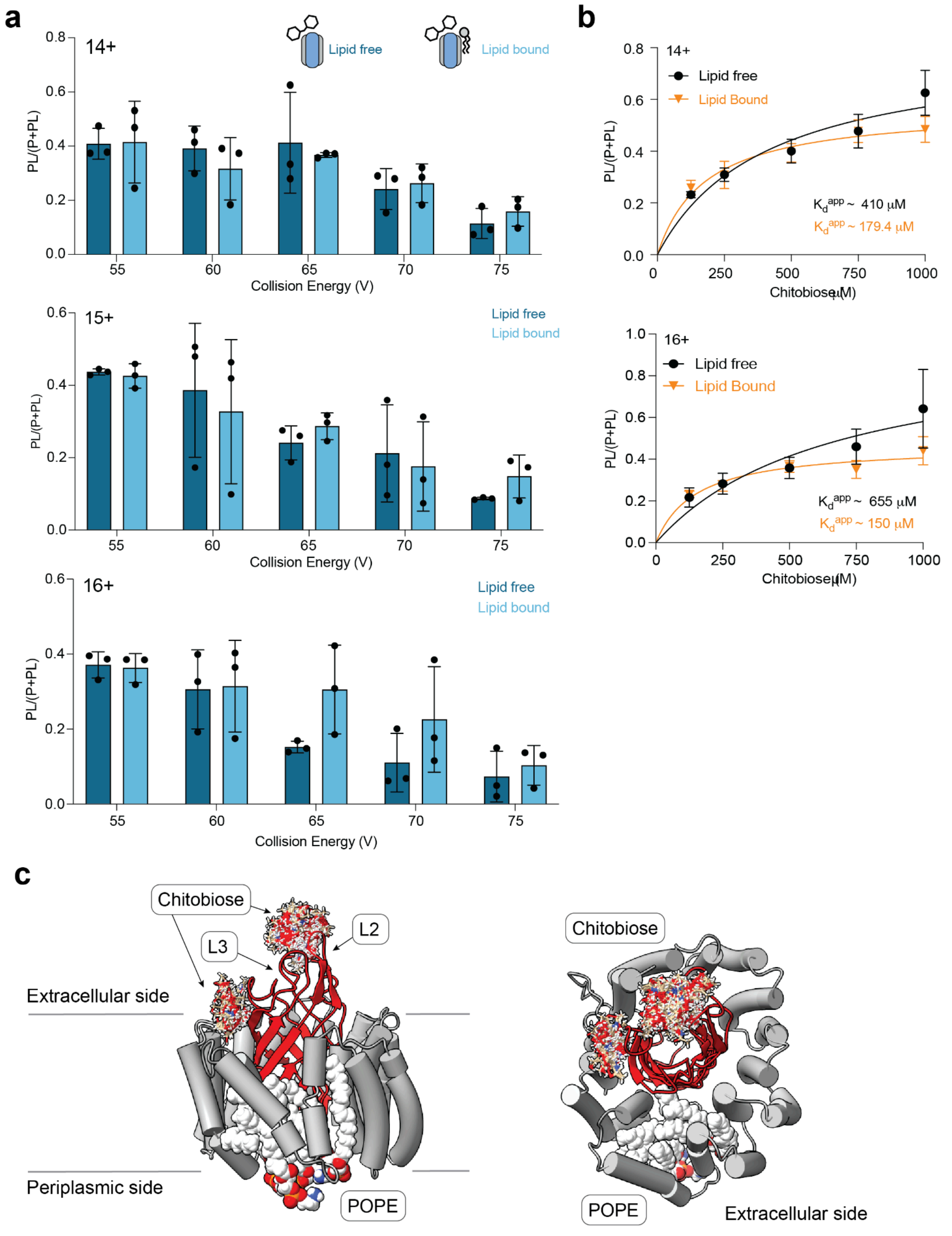


***Figure S5.*** *(a) Dissociation of bound chitobiose from lipid-free and PE-bound WRAP-OmpA as a function of activation (ion trap voltage). Data are shown as average and standard deviation (n = 3). (b) Plots of the ratio of PL to P + PL for WRAP-OmpA and WRAP-OmpA-PE binding to chitobiose for the 14+ and 16+ charge states of the complex. Error bars indicate average and standard deviation (n = 4). (c) SwissDock model[10] of chitobiose binding to the WRAP-OmpA-4xPE complex. The lipid-binding residues are shown as sticks, possible chitobiose orientations are rendered as an overlay of the top 20 conformers. Chitobiose occupies the previously identified extracellular loops 2 and 3 as well as a secondary binding site on the side of the central barrel, whereas PE interacts with the periplasm-facing area on the opposite side of the protein.*
